# Linking null models to hypotheses to improve nestedness analysis

**DOI:** 10.1101/2021.04.05.438470

**Authors:** Rafael Barros Pereira Pinheiro, Carsten F. Dormann, Gabriel Moreira Félix Ferreira, Marco Aurelio Ribeiro Mello

**Affiliations:** Graduate School in Ecology, Conservation and Wildlife Management, Federal University of Minas Gerais, Brazil; Department of Biometry and Environmental System Analysis, University of Freiburg, Germany; Graduate School in Ecology, University of Campinas, Brazil; Department of Ecology, University of São Paulo, Brazil

**Keywords:** community ecology, interaction matrices, metacommunity, network science, network topology, nestedness, null model analysis, species interactions

## Abstract

1. Nestedness is a common pattern in metacommunities and interaction networks, whose causes are still subject to investigation. Nestedness inference is challenging because, beyond calculating an index, we need to compare observed and expected values using null models. There are many different null models and the choice between them strongly affects test outcomes. Furthermore, there is no theoretical basis to guide this choice.
2. Here, we propose a different look at the meaning of nestedness that improves our understanding of its causes and directly links null models to hypotheses. The nestedness of a matrix is the combination of marginal sum inequality and high overlap. The higher the overlap, the more predictable the cell values by marginal sums. Therefore, the only way a process can increase nestedness is by promoting marginal sum inequalities, without concomitantly introducing preferences. We propose a combination of null models to test for different topological hypotheses. In these null models, interactions are randomized following a probability matrix. The probability matrix of the equiprobable model excludes marginal sum inequalities and consequently all nestedness-generating mechanisms, providing a distribution of expected values for nestedness significance tests. The probability matrix of the proportional model, by conserving marginal sums and maximizing overlap, conserves nestedness-generating mechanisms and excludes nestedness-disrupting mechanisms, thereby yielding the expected nestedness values for fully nested matrices.
3. We advance a new protocol to test for nestedness in binary and weighted matrices, in which those two null models represent different topological hypotheses. Our protocol helps to tell apart cases in which nestedness is different from expected by chance and cases in which nestedness represents the best topological archetype. We also evaluate the efficiency of applying this protocol using some common nestedness indices. Analyses based on weighted data proved to be highly efficient and informative, while binary analyses induced severe distortions. Finally, we empirically illustrate our approach using a plant-pollinator network.
4. Through a shift of perspective, our approach reconciliates contradictions in null model analysis and provides a guideline for nestedness interpretation.

## 1. INTRODUCTION

The concept of nestedness originated in the study of island biotas. Nestedness was first defined as a biogeographical pattern in which faunas of species-poorer islands within an archipelago constitute a subset of the faunas of species-richer islands (Patterson & Atmar, 1986). These patterns had been recognized earlier by biogeographers (Darlington, 1957). Nevertheless, it was only after the influential work of Diamond (1975) that the structure of metacommunities, using species-by-sites matrices, turned into a major focus of ecological studies, boosting the search for nestedness. In the following decades nestedness in species occurrences has been found in islands (Wright & Reeves, 1992), gradients (Harrison et al., 1992), mountain ranges (Cutler, 1991), and several other contexts (Wright et al., 1997). Later, the observation of nestedness in species interaction networks (Bascompte et al., 2003) highlighted it as a very common pattern in ecological systems.

Despite the ubiquity of nestedness, its drivers in ecological systems remain disputed (Valverde et al., 2018; Pinheiro et al., 2019). Nestedness in species occurrences is traditionally explained as the result of ordered selective colonization (Darlington, 1957; Patterson & Atmar, 1986) or extinction (Cutler, 1991) in local communities. The resulting order could arise, for instance, as the combination of differential colonization abilities (or extinction vulnerabilities) by species and differential probabilities of colonization (or extinction) depending on island size and isolation (Cutler, 1994). In contrast, nestedness in species interactions is often related to unequal probabilities of species to interact with others (Ulrich & Almeida-Neto, 2012), resulting, for instance, from unequal abundances (Vázquez et al., 2007; Krishna et al., 2008).

To investigate the drivers of nestedness, it is first necessary to operationalize the concept and measure how nested real-word networks are. To do so, networks are usually represented as bipartite interaction matrices (i.e. incidence matrices), where rows and columns represent the nodes (e.g. pollinators and plants) and cells represent the links between them (e.g. floral visits). There are several nestedness indices based on different calculations and definitions (Ulrich et al., 2009). A notable advancement was the development of nestedness indices for weighted matrices (Almeida-Neto & Ulrich, 2011). In real-world networks, interactions between nodes are not all equivalent, but vary in frequency or intensity. For instance, links in a pollination network might be weighted by the frequency of pollination events observed for each plant-pollinator combination (Watts et al., 2016). Weighted matrices are more informative and, often, reveal patterns that are hidden in their binary counterparts (Banašek-Richter et al., 2004; Scotti et al., 2007). Nevertheless, in the end, the choice between weighted and binary matrices depends on the question under investigation (Fründ et al., 2016; Jordano, 2016).

However, before invoking causal mechanisms, it is important to test for the significance of the observed nestedness value, since even random matrices will present some degree of nestedness. To infer significance, the observed nestedness values are usually compared with the distribution of values calculated for randomized matrices generated by null models (Ulrich et al., 2009). Null models are randomization procedures that conserve more or less inclusive subsets of the original matrix properties. Therefore, null models are actually null hypothesis about the mechanisms needed to produce an specific pattern (Gotelli & Graves, 1996).

Algorithms to generate null models differ markedly. Although the choice of null models strongly affects test outcome (Ulrich & Gotelli, 2007), there is no theoretical basis to guide this decision in nestedness analysis. Null models that only conserve matrix dimensions and total sum are considered less rigorous, but more sensitive, to nestedness tests, than null models that conserve matrix marginal sums. In addition, as algorithms are not linked to explicit hypotheses, this choice is often regarded as a trade-off between type I and type II statistical errors (Ulrich & Gotelli, 2007; Strona et al., 2018). This is a questionable interpretation, as different null models likely exclude the effects of different ecological mechanisms, and hence might not be appropriate to test the same null hypothesis. Solving this dilemma requires a better understanding of how the mechanisms excluded or conserved by each null model relate to the emergence of nestedness, so that we can link each null model to a specific topological hypothesis.

In the present study, we propose a novel perspective on nestedness in interaction matrices and a protocol to operationalize it. We believe that both the way in which observed nestedness values are compared with null models and the interpretation of these comparisons can be improved. In several steps, we firstly define nestedness in a matrix as a combination of marginal sum inequalities and overlap. Secondly, we show that overlap measures how well marginal sums predict cell values. Thirdly, we argue that two commonly used null model algorithms are neither alternative nor conflicting, but complementary, as they simulate the exclusion of different mechanisms and, thus, test different hypotheses. The equiprobable null model excludes all nestedness-generating mechanisms and is appropriate for significance tests. The proportional model conserves all nestedness-generating mechanisms and excludes nestedness-disrupting mechanisms, so it can be used to test whether the matrix significantly deviates from a fully nested topology. Fourthly, we evaluate the efficiency of this protocol in combination to common nestedness indices using both binary and weighted matrices. Lastly, we illustrate these new procedures using a pollination network.

## 2. MATERIALS AND METHODS

### 2.1. A NEW PERSPECTIVE ON THE MEANING OF NESTEDNESS

While we focus our examples on species interaction networks, our rationale may be extended to other bipartite networks, both ecological (e.g. species vs. sites) and non-ecological (e.g. actor vs. movies). Thus, for the sake of generality, here we use common terms for the study of interaction matrices and networks.

#### Orders of information in an interaction matrix

Matrix information can be partitioned into three levels (Dormann et al., 2017). First-order information comprises the most general features of the interaction matrix: its dimensions (number of rows and columns) and the sum of all cell values (hereafter, total sum). Second-order information represent the specific properties of each row and column: the marginal sums. Third-order information is the actual cell values. In Fig. 1, we illustrate these orders of information using a plant-animal interaction matrix.

**FIGURE 1.**
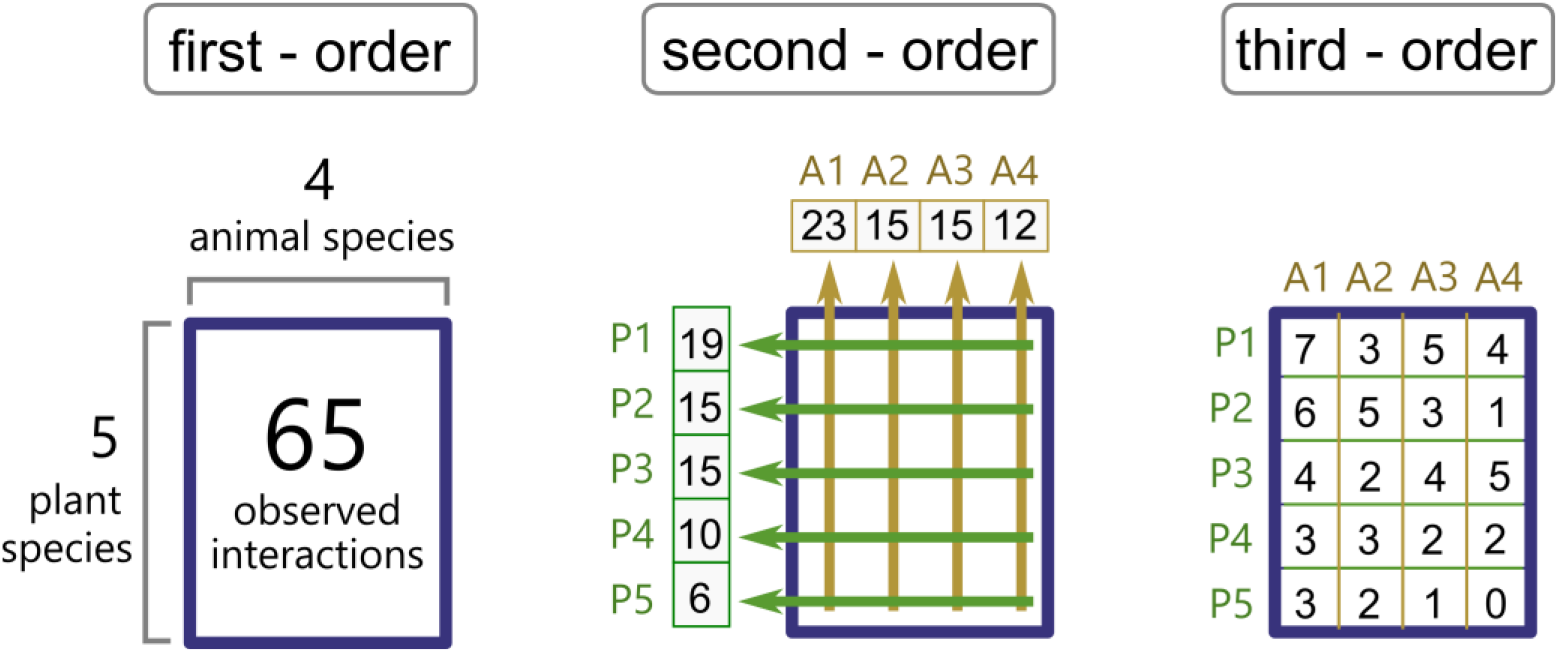
Orders of information in a weighted plant-animal interaction network. First-order refers to the general aspects of the matrix: the species richness on each side of the interaction (dimensions) and the total frequency of interactions observed (total sum). Second-order refers to specific properties of the rows and columns: the frequency of interactions made by each plant and animal species (marginal sums). Third-order refers to the pairwise interactions between rows and columns: the frequency of interactions between each plant-animal pair (cell values).

#### Definition of nestedness

The concept of nestedness was first proposed as a pattern of species occurrence, latter generalized to binary bipartite networks, and only recently applied to weighted bipartite networks. Thus, it is not trivial to provide a general definition of nestedness for all kinds of matrices. Below, we provide a verbal definition, and additionally, a graphical definition, so it is easier to understand and operationalize the concepts.

Bipartite interaction networks are usually represented as matrices, within which the two classes of nodes are represented by rows and columns. To calculate nestedness, we always compare rows with rows, and columns with columns. Binary nestedness is, the tendency of rows with lower marginal sums to be linked (cell value = 1) with a subset of the columns linked to the rows with higher marginal sums; and tendency of columns with lower marginal sums to be linked (cell value = 1) with a subset of the rows linked to the columns with higher marginal sums. For weighted nestedness, beyond the identity of linked nodes, we may account for link weights (cell values). Thus, weighted nestedness is the tendency of rows with lower marginal sums to have weaker links (lower cell values) with the same set or a subset of the columns linked to the rows with higher marginal sums; and the tendency of columns with lower marginal sums to have weaker links (lower cell values) with the same set or a subset of the rows linked to the columns with higher marginal sums.

To visualize nestedness we must organize rows and columns by decreasing marginal sums. In a binary matrix, nestedness is a tendency of 1s to be in the top-left, while 0s are in the bottom-right corner of the matrix. In a weighted matrix, nestedness is a tendency of decreasing cell values from the top-left to the bottom-right corner (Fig. 2a).

**FIGURE 2.**
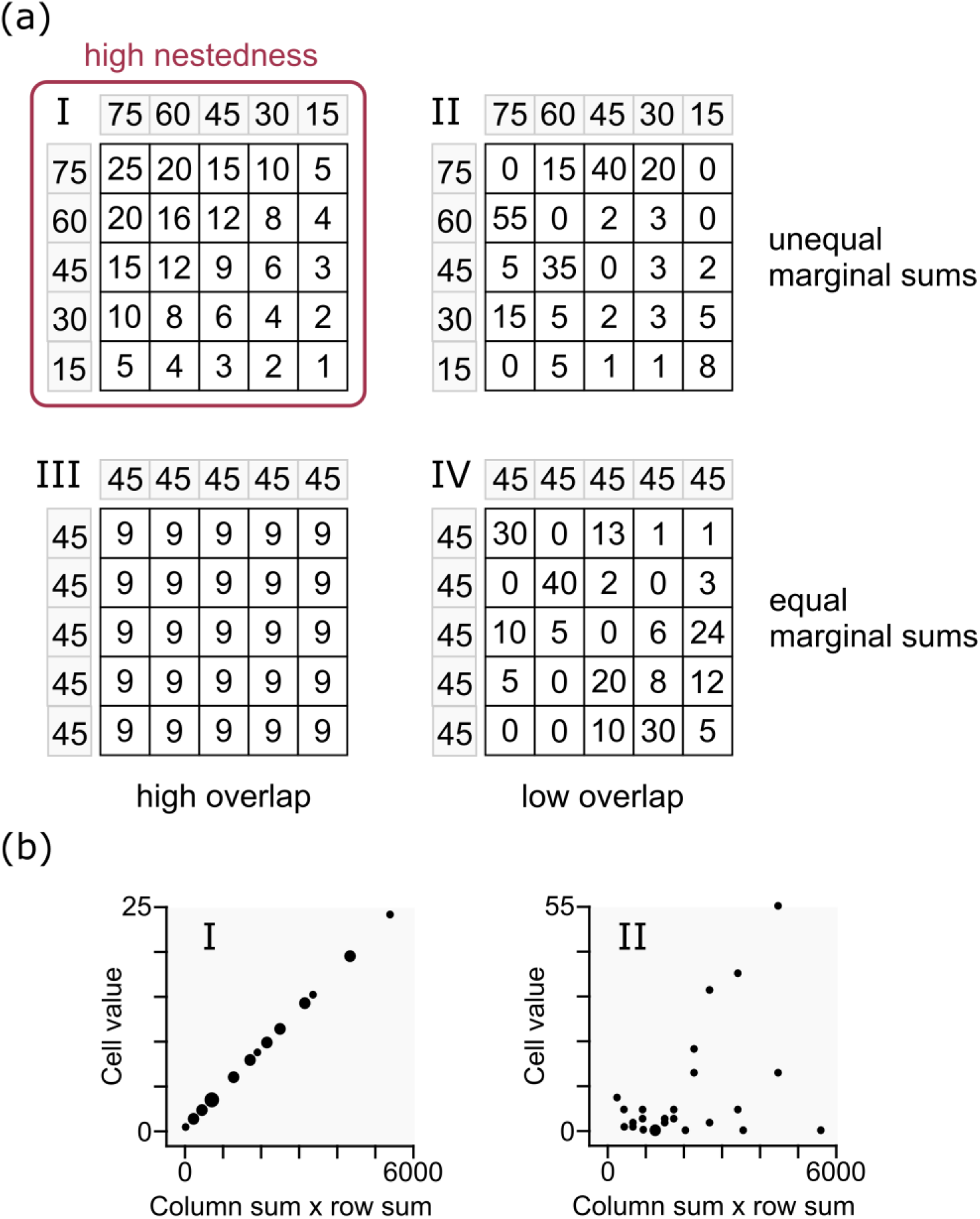
(a) Nestedness is a combination of high overlap and marginal sums inequality. Weighted overlap is the tendency of rows to establish links with the same columns and to distribute weights among these columns in similar proportions, and the tendency of columns to establish links with the same rows and to distribute weights among these rows in similar proportions. Let us imagine four matrices: two with unequal marginal sums (I and II, see the marginal sums in the grey cells), two with equal marginal sums (III and IV); two with high link overlap (I and III), two with low link overlap (II and IV). In a matrix with unequal marginal sums, high overlap results in nestedness. Here, we focus on weighted matrices, as similar explanations for binary matrices were already provided by other studies (Almeida-Neto et al., 2008; Ulrich & Almeida-Neto, 2012). (b) Overlap in a matrix may alternatively be understood as a measure of how much each cell value adjust to the expected given the outer product of the marginal sums of its respective row and column (second-order information). Notice that cell values increase monotonically with the outer product of their marginal sums in the nested matrix (I) but not in the matrix with low overlap (II).

Some authors argue that weighted nestedness should require binary marginal sum inequalities (decreasing degrees), so that weighted nestedness in a matrix is constrained by binary nestedness (Almeida-Neto & Ulrich, 2011). This definition aims at a better match between the concepts of nestedness and ecological specialization. Despite making sense for the study of ecological networks, this definition yields a mixed binary/weighted topology rather than a purely weighted topology. Here, we opt to address separately the binary and the weighted structures of a matrix, which we believe leads to a clearer and more general interpretation.

Binary overlap is the tendency of different rows to link to a similar set of columns, and vice versa. The core of any nestedness index is to measure the overlap between nodes with different marginal sums. In other words, to measure how often rows with lower marginal sums are linked to the same columns as rows with higher marginal sums; and how often columns with lower marginal sums are linked to the same rows as columns with higher marginal sums. Weighted overlap indices account for not only the identity of linked nodes, but whether nodes of the same class distribute weights among their links similarly. The opposite of overlap, a tendency of different rows to interact with different columns is often called *turnover, specialization*, or *preferences*. Here, we opt for *preferences*, but notice that this term here refers strictly to a matrix pattern, not necessarily matching the homonymous ecological concept.

As pointed out by previous studies (Baselga, 2012; Ulrich & Almeida-Neto, 2012), the original concept of nestedness also assumes a gradient in marginal sums. This realization prompted the development of nestedness indices explicitly demanding marginal sum inequalities, such as NODF (Nestedness based on Overlap and Decreasing Fill) (Almeida-Neto et al., 2008). Nestedness is thus a combination of two features: high overlap and unequal marginal sums (Figure 2a).

#### The meaning of overlap

Here, we propose an alternative perspective on the meaning of overlap with implications for the understanding of nestedness. Imagine that the nested weighted matrix (I) in Fig. 2a represents the observed frequency of interactions between five pollinator species (columns: C1, C2, C3, C4, and C5) and five plant species (rows: R1, R2, R3, R4, and R5). A high overlap of interactions between plant species means that the relative frequency of interactions with each pollinator species is similar for all plants and, thus, determined by the total interactions of each pollinator (column sums). For instance, as the pollinator C1 interacts more frequently with the plant R4 than does pollinator C4 does, we would expect it to also interact more frequently with any other plant species than C4 does. Applying the same logic to the transpose, we conclude that the frequency of interactions between each pollinator-plant pair in a nested matrix can be well predicted by the total frequency of interactions of each pollinator and each plant species (marginal sums) (Fig. 2b). This same idea has been previously applied to binary matrices, Ryti & Gilpin (1987) used the R² of a logistic regression of occurrences as a measure of orderedness (overlap).

This is the first key insight of our study: the higher the overlap in a matrix, the higher the predictive power of the second-order information (marginal sums) for cell values. However, when all marginal sums are equal, they provide no additional information about cell-values compared to a prediction based only on the first-order information (dimensions and total sum). In a fully homogeneous matrix (Fig. 2a III), by knowing the marginal sums, we can perfectly predict all cell values, but we achieve the same result by just averaging cell values. Thus, the second key insight of our study: for a given overlap, the higher the inequalities in the marginal sums, the higher the differences between a prediction about cell values informed by second-order information and a prediction only based on first-order information.

Putting these two insights together, we find that nestedness is directly related to how much information about cell values we gain by knowing, not only the first-order, but also the second-order properties of a network. In a matrix with high nestedness, it is a much better prediction to assume that each cell value is proportional to the outer product of row and column sums than to guess that all cell values are the same. This understanding is not opposed or divergent from the currently most accepted nestedness definition (Almeida-Neto et al., 2008; Ulrich & Almeida-Neto, 2012), but a logical implication from it.

#### Nestedness-generating and nestedness-disrupting mechanisms

It follows from the above discussion that the only way a mechanism can generate nestedness in a matrix is by promoting marginal sum inequalities without decreasing overlap. Therefore, nestedness-generating mechanisms must drive the second-order information of a matrix. On the other hand, mechanisms that introduce preferences (i.e. decreases overlap) on the matrix cause nestedness disruption. Thus, nestedness-disrupting mechanisms causes deviations in cell values (third-order information) from the expected values given the marginal sums (second-order information).

### 2.2. LINKING NULL MODELS TO HYPOTHESES

#### Equiprobable null model: testing nestedness significance

To infer the significance of nestedness in an interaction matrix, we must first state our expectations using a null model. The defining characteristic of a null model is that *it excludes the mechanism being tested* (Gotelli, 2001). Considering our definitions, to exclude all nestedness-generating mechanisms, such a null model should discard second- and third-order information of the matrix, so that the two defining elements of nestedness - inequality of marginal sums and overlap – emerge randomly. Thus, this null model must only conserves first-order information: matrix dimensions and total sum, hereafter called “equiprobable null model” (Ulrich et al., 2009). This model corresponds to a fully weighted version of the Erdös-Rényi model (Erdös & Rényi, 1959) or the null model 1 of Bascompte et al. (2003), in which the total sum of interactions rather than connectance is conserved.

In the equiprobable null model, interactions are distributed on randomized matrices with the same dimensions as the observed matrix, and cells have equal probability to receive an interaction. Randomized matrices, thus, are realizations from a probability matrix whose nestedness is always zero (Fig. 3). An equiprobable null model provides the nestedness distribution expected for matrices with random topologies, excluding all nestedness-generating mechanisms, and yields a benchmark for significance tests of nestedness.

**FIGURE 3.**
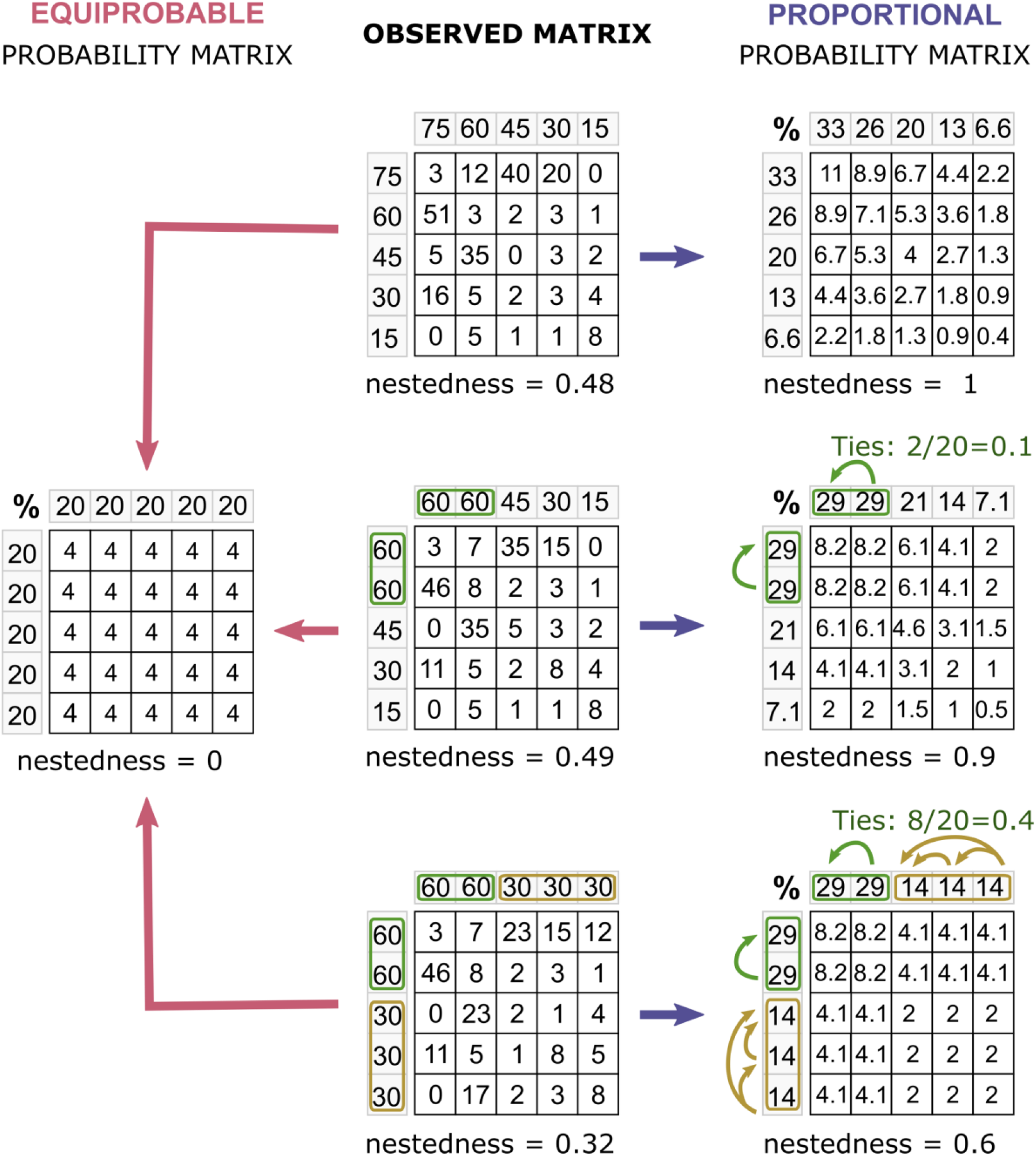
To test nestedness, the observed value is usually compared to the distribution of values from randomized matrices generated by null models. Here, we illustrate the probability matrices of two null models: equiprobable and proportional. In both models, the total sum of observed interactions is distributed in randomized matrices with equal dimensions following these probabilities. In the equiprobable null model, any cell has the same probability of receiving an interaction and, thus, the nestedness of the probability matrix is always zero. In the proportional null model, on the contrary, probabilities are defined by the outer product of the relative marginal sums, so that they are maximally nested for a given marginal sum distribution. In these 5×5 matrices, for instance, there are 20 pairwise comparisons between nodes (10 between rows and 10 between columns). When there are no marginal sum ties in the observed matrix, nestedness of the probability matrix in the proportional null model is always 1. Otherwise, nestedness decreases in the exact proportion of ties, representing the maximum achievable for such marginal sums. Nestedness was measured with the WNODA index (Pinheiro et al., 2019).

#### Proportional null model: testing for a fully nested topology

After inferring that a given network presents significant nestedness compared to a random topology, the logical next question is whether it significantly deviates from a fully nested topology. In this case, we must state the expectations through a null model that conserves all nestedness-generating mechanisms and discards all nestedness-disrupting mechanisms. Accounting for our definitions, such a null model must conserve first- and second-order information of the randomized matrices, so that marginal sum inequalities are kept. Then, it must randomize third-order information to discard preferences.

The appropriate algorithm for this task is one in which the probability of each cell in the randomized matrix to receive a unit of weight is proportional to the respective marginal sums of the observed matrix, hereafter, called proportional null model (Gotelli, 2000). Probabilities in a proportional model are maximally nested for a given marginal sum distribution: the overlap of probabilities is maximum and, thus, nestedness in the probability matrix is only decreased by the proportion of marginal sum ties (Fig. 3). This model corresponds to a fully weighted version of the null model 2 of Bascompte et al. (2003).

The proportional model provides the expected nestedness distribution for matrices whose topology is only shaped by nestedness-generating mechanisms. By comparing the observed nestedness with the distribution generated by a proportional model, we can test whether the observed matrix is a likely realization from a maximally nested probability matrix. This understanding conflicts with the common use of proportional models as a null distribution to test nestedness significance. This model should be considered the upper distribution of expectation for nestedness, rather than the lower distribution. Otherwise, as earlier suggested by Wright and Reeves (1992), the null model itself hides the pattern that we attempt to detect. Indeed, when the proportional or other weighted null models that conserve marginal sums is applied as null distribution, network are almost universally detected as non-nested or anti-nested (Joppa et al., 2010; Staniczenko et al., 2013).

Notice that both the equiprobable and the proportional models we discuss here do not have additional binary constraints, such as maintaining connectance (Vázquez *et al*., 2007), as this is not a first-neither a second-order property of weighted matrices. This mixed use of binary and weighted information is undesirable for our approach, as it cannot be linked to any specific hypothesis about nestedness.

#### A new protocol for testing nestedness

Nestedness is a quantitative network property that, even when significant, might vary broadly in intensity. Although nestedness can and should be understood as a continuum, there are often pragmatic and philosophical reasons to classify networks into topological categories (Lewinsohn *et al*., 2006). Here, we discourage the use of nestedness significance to delimit a topological category. Nestedness significance compared to an equiprobable null model just means that the observed pattern is unlike to emerge without some mechanism that affects marginal sum distributions, but it does not rule out the action of other network-shaping mechanisms. When significance is used as the unique criterium, a variety of structures that are not sufficiently coherent are likely merged into a single category. On the other hand, a proportional null model is composed of matrices only shaped by nestedness-generating mechanisms. In a mechanistic perspective, matrices of a proportional null model are as nested as a nested matrix could be, given its dimensions and marginal sums, and, thus, they are ideal types to represent a nested topology (Fig. 4).

**FIGURE 4.**
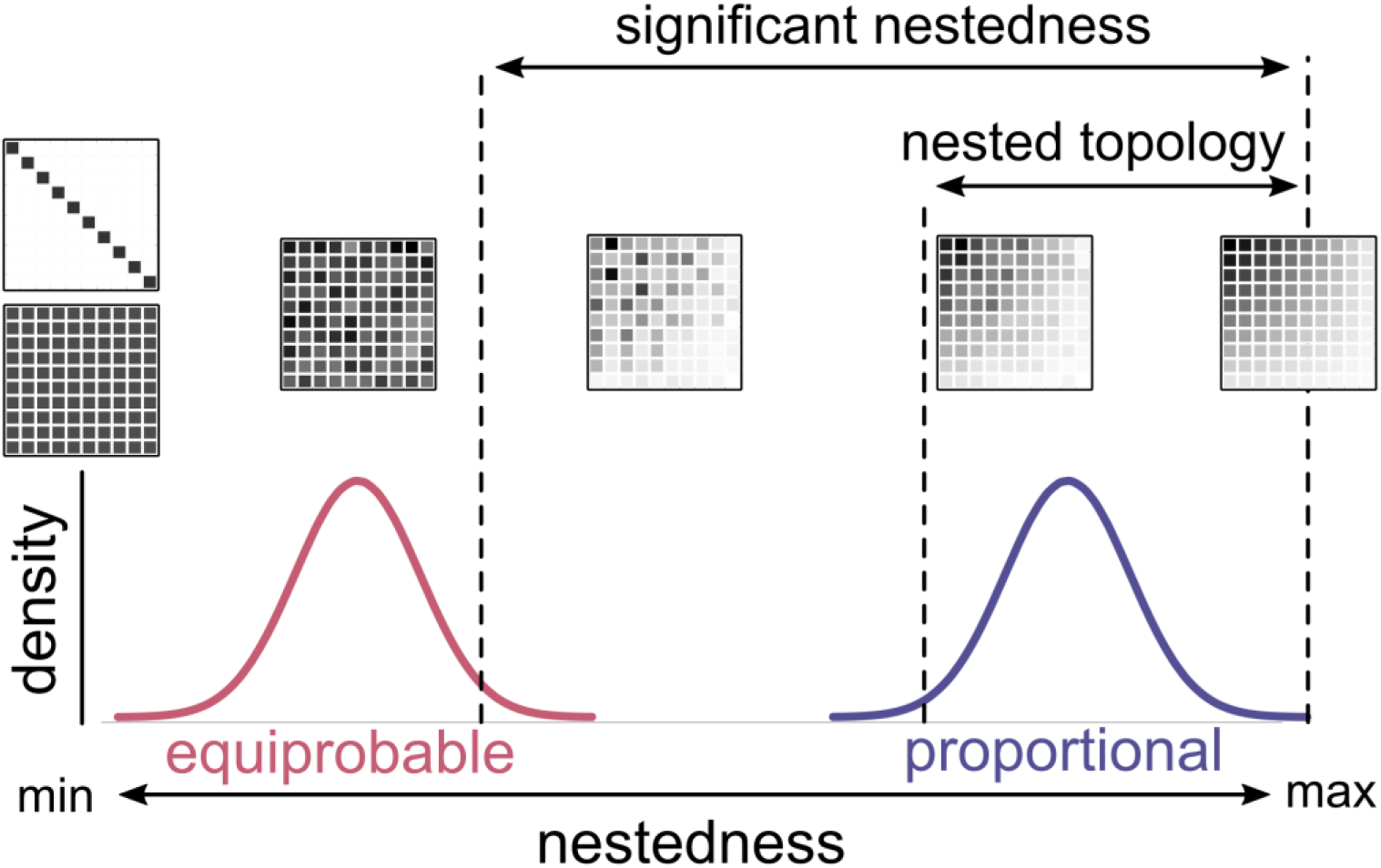
A new protocol for the use of null models in tests of nestedness. The equiprobable null model excludes all nestedness-generating mechanisms and represents a truly random topology. Nestedness significance is checked by the comparison of the observed value with the distribution of values generated by the equiprobable null model. The proportional null model is composed of matrices whose interactions derive from maximally nested probabilities. It conserves the effects of nestedness-generating mechanisms and excludes the effects of nestedness-disrupting mechanisms, resulting in a nested topology. Observed nestedness values should be interpretated in comparison to the distributions generated by these null models, considering what mechanisms each of them conserves or discards, and what topology each of them shows. Nevertheless, when the aim is to classify networks into topological categories, we recommend two criteria to delimit a fully nested topology: first, nestedness must be significantly higher than the equiprobable model; second, nestedness must not be significantly lower than expected by the proportional model.

Accounting for this new interpretation of null models in nestedness analysis, we suggest a new protocol to test for nestedness. The steps of this procedure are:

1. Calculate nestedness for the observed matrix.
2. Build an equiprobable null model.
3. Infer nestedness significance using randomized values generated with the equiprobable model as the null distribution.
4. If nestedness is significant, build a proportional null model.
5. Use the proportional null model to generate the expected distribution of nestedness for matrices with a nested topology. Thereby, check whether the observed nestedness is coherent with the distribution of values generated with the proportional model. If the observed nestedness does not significantly differ from the proportional null model, the matrix’s topology can be classified as fully nested.
6. Plot the observed value and null model distributions of nestedness, prompting a continuous analysis of where a given matrix falls within the possibility space illustrated in Fig. 4.

R functions for using this procedure are available on GitHub (https://github.com/pinheirorbp/nestedness).

### 2.3. REMAINING CHALLENGES

In real-world systems, interactions almost always have a weighted nature (e.g. frequencies and abundances). However, due to data collection limitations, networks are often studied using binary matrices (Dormann & Blüthgen, 2017). When based on binary data, marginal sum distributions are highly contingent on connectance (Poisot & Gravel, 2014), and so they are poor proxies for the respective weighted marginal sums, especially for matrices with high connectance (Fig. S1a). As a result, binary proportional models often cannot conserve the marginal sum distribution on the randomized matrix (Wright et al., 1997). In addition, without weighted information, null models can only conserve connectance. Connectance, however, is not a direct proxy for the first-order information of the weighted matrices, as it is also highly affected by marginal sums and preferences (Fig. S1b).

Thus, when connectance is kept, null models cannot be fully distinguished by the orders of information conserved or discarded. Such distortions are likely to cause severe misestimation of nestedness in binary null models.

### 2.4. EFFICIENCY ANALYSIS

We used simulated matrices to analyze the efficiency of our protocol in combination with the most used binary nestedness index, NODF, and two weighted versions of it, Weighted Nestedness based on Decreasing Fill (WNODF) (Almeida-Neto & Ulrich, 2011) and Weighted Nestedness based on Decreasing Abundance (WNODA) (Pinheiro et al., 2019), in order to distinguish between matrices with random (equiprobable null model) and nested (proportional null model) topologies. WNODF is a mixed binary-weighted index, as it measures weighted overlap, but requires decreasing binary marginal sums, while WNODA is a fully weighted index (see Pinheiro et al., 2019).

First, we produced probability matrices with dimensions: 5 × 5, 10 × 10, and 20 × 20, based on three different marginal probabilities: constant, linear decrease, and log-normal. These represent a gradient between an equiprobable null model and a proportional null model based on a more skewed marginal sum distribution. Then, following cell probabilities, we distributed interactions in simulated matrices with different sampling intensities (i.e. the total frequency of interactions distributed in each matrix): 50, 100, 200, 400, 800, 1600, 3200, and 6400 (Fig. S2). For each unique setup we produced 10,000 matrices. For each matrix we calculated NODF, WNODF and WNODA and graphically inspected the capacity of separating results between the equiprobable and the proportional (linear decrease and lognormal) models. Indices were calculated using the bipartite package for R (Dormann et al., 2008).

As binary matrices do not hold weighted information, we cannot fix the true total sum of weights on the null models, only connectance. For a more realistic assessment of NODF, as a binary index, we generated binary matrices from the probabilities with fixed connectance of 0.3, 0.5, 0.7, 0.9, 0.92, 0.94, 0.96, and 0.98. Moreover, in analysis of binary matrices, we usually do not know the original weighted marginal sums, only its binary respective. To inspect the distortion caused by this approximation, from each matrix produced with fixed connectance, we produced an additional randomized matrix with the same connectance using a proportional algorithm based on binary marginal sums (null model 2 of Bascompte *et al*., 2003).

An issue with any randomizing algorithm based on probabilities is that it may produce matrices with empty rows or columns. In general, these are not likely to bias network analysis (Gotelli, 2000). Here, for the weighted matrices we kept all matrices. However, reducing matrix dimension modifies its connectance. Thus, when producing binary matrices, we discarded all the matrices with reduced dimensions.

### 2.5. CASE STUDY

To illustrate our novel perspective we used a weighted plant-pollinator network, originally described by Kaiser-Bunbury *et al*. (2010) and available in the Web of Life database (www.web-of-life.es, network: M_PL_060_17). The network includes 52 nodes, which represent 17 plant species and 35 pollinator species. Weights represent the frequency of floral visits observed between each pollinator and each plant species (total of 444 visits). This network was arbitrarily chosen from the database, without any preliminary analysis.

We produced an equiprobable and a proportional null model, with 10,000 matrices each, and calculated WNODA for each matrix. Similarly, we analysed the binary structure of the network using NODF. Then, we produced a binary equiprobable null model in which connectance, rather than total sum of weights, was conserved in the randomized matrices. For last, we produced a proportional model based on binary information: binary marginal sums were used to calculate probabilities and connectance was conserved in the randomized matrices, to test whether it would result in a distortion of NODF values compared to the proportional model based on weighted data.

## 3. RESULTS

### Efficiency analysis

We found that, using our protocol, WNODA has an outstanding efficiency in discriminating between weighted matrices with random and nested topologies (Fig. 5A, Appendix S1). As expected, for all indices, nested matrices with a more skewed marginal sum distribution (log-normal) are more easily distinguished from the equiprobable null model than nested matrices with more similar marginal sums (linear-decrease). The efficiency of WNODA increases with increasing sampling effort, and values tends to stabilization. NODF and WNODF, however, due to the use of binary information (binary overlap for NODF and decreasing fill for both), are more efficient with intermediary sampling and are not appropriate for using with highly filled matrices.

**FIGURE 5.**
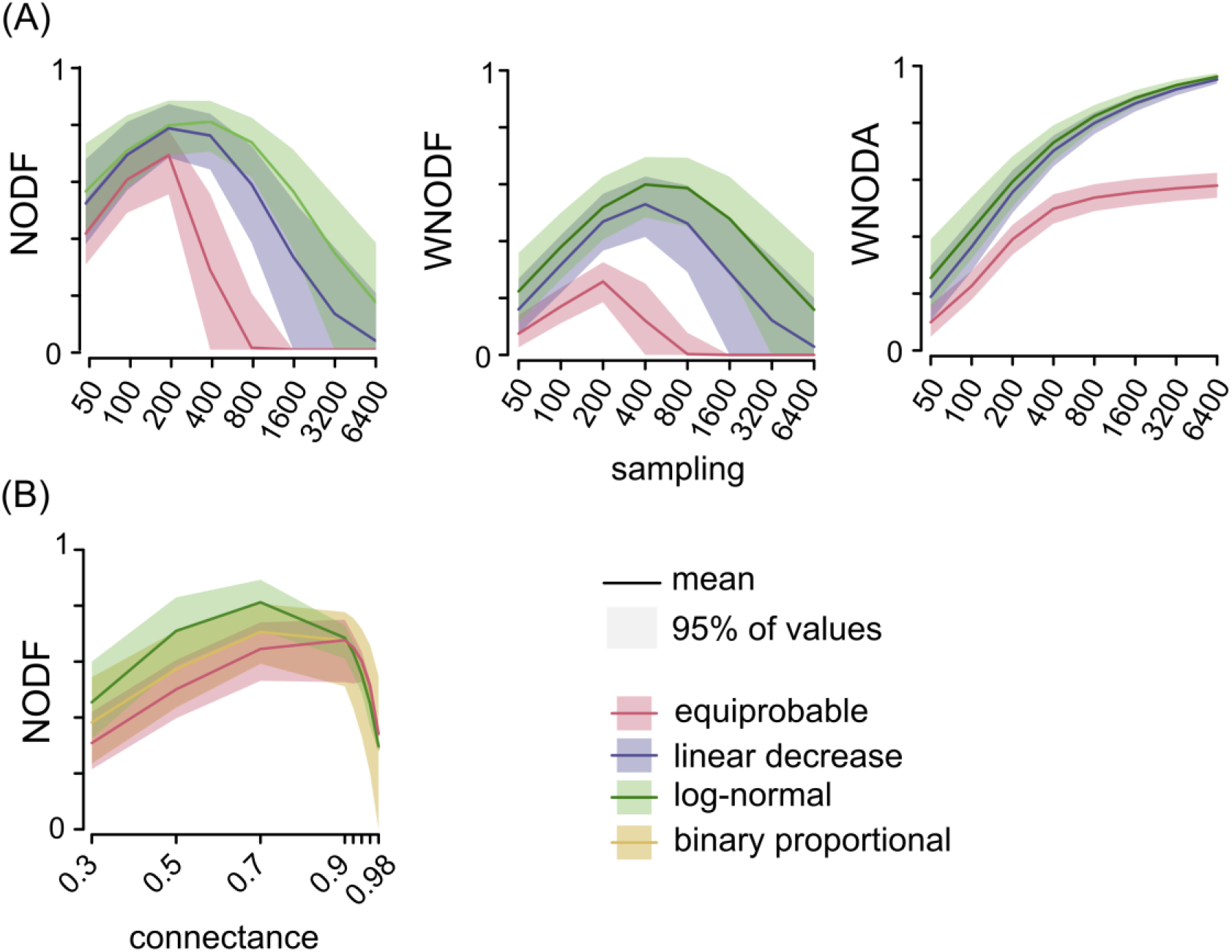
Ability of indices to distinguish between nested matrices (cell values are determined by marginal sums: linear decrease and log-normal) and random matrices (cell values are randomly distributed: equiprobable) following our protocol. The larger the separation between the distributions of random and nested matrices, the more efficient the analysis. In (A), indices are applied to weighted matrices produced with different sampling intensities. WNODA shows an outstanding efficiency compared to the other indices. In (B), NODF is applied to binary matrices produced with different connectances. As binary matrices do not provide information on weighted marginal sums, we produced a proportional model based on the binary marginal sums of the matrices generated with the log-normal probabilities. Ideally, values for the binary proportional model (yellow) would match values for the matrices based on log-normal probabilities (green), but we found that they are much lower and closer to the equiprobable values (red). Only results from the 10 × 10 matrices are included in the figure, see Appendix S1 for other matrix dimensions.

Weighted information on the matrix marginal sums is not available in binary matrices, and the use of binary information to estimate probabilities on the proportional model resulted in large distortions. NODF values for matrices of the proportional null model based on the binary marginal sums were much lower than values for the matrices produced from weighted information (Fig. 5B).

### Case study

The empirical plant-pollinator network showed significant nestedness (WNODA = 0.18, comparison against the equiprobable model: Z = 2.73, p = 0.006), but cannot be classified as presenting a nested topology (comparison against the proportional model: Z = -6.73, p < 0.001; Fig. 6: Weighted nestedness). The binary index (NODF) leads to similar conclusions (NODF: 0.30; comparison against the equiprobable model: Z = 6.61, p < 0.001; comparison against the proportional model: Z = -4.87, p < 0.001). The distribution produced by the binary equiprobable null model (Fig. 6: Binary nestedness) was almost identical to the distribution produced by the weighted equiprobable null model, resulting in similar results (Z = 6.70, p < 0.001). However, when the proportional model was based in binary information, there was a large underestimation of nestedness in the randomized matrices, and because of that, the observed nestedness is seemingly too high compared to the proportional model (Z = 3.31, p < 0.001).

**FIGURE 6.**
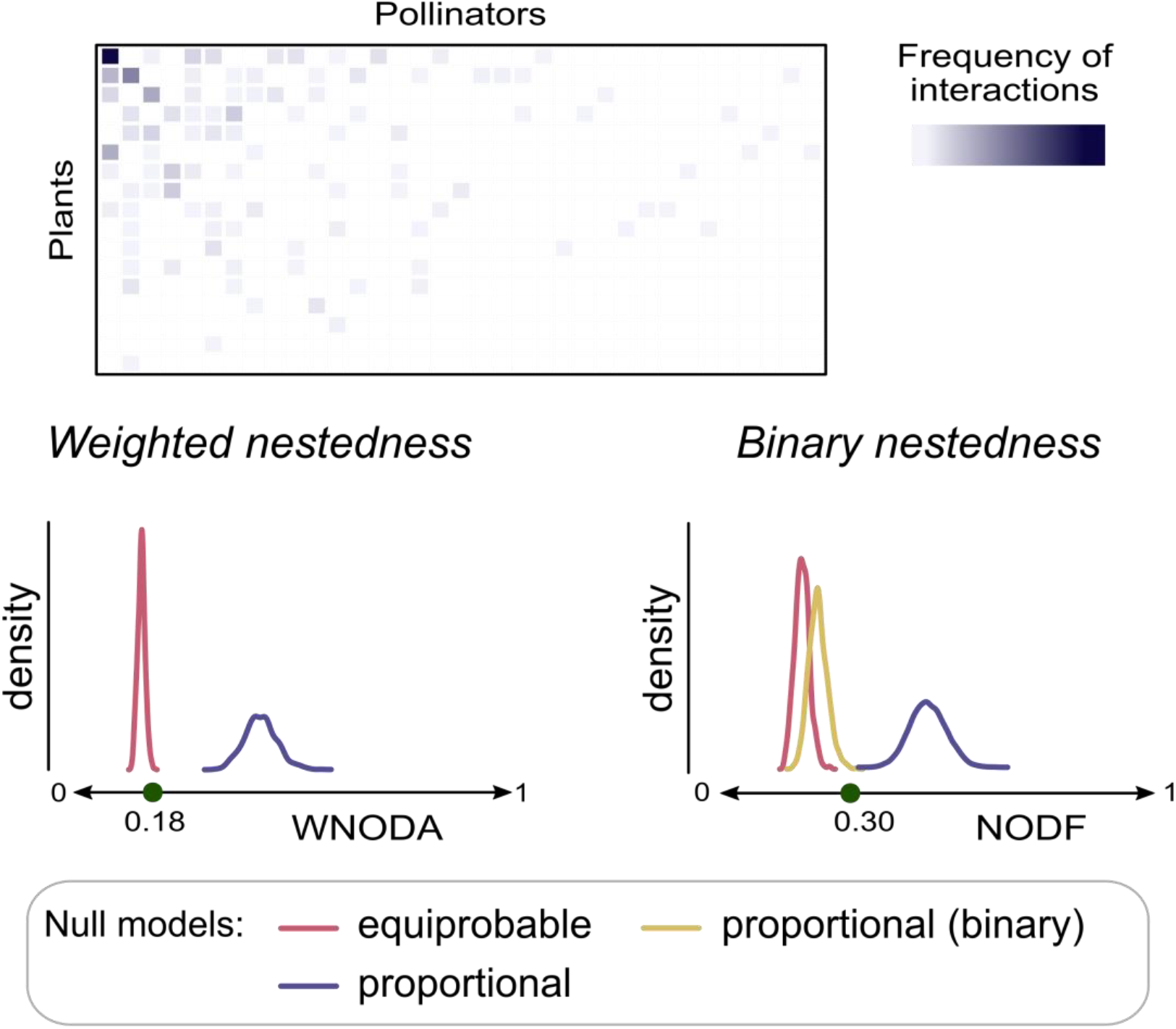
Case study using a weighted empirical plant-pollinator network (Kaiser-Bunbury et al., 2010) to illustrate the new protocol. *Weighted nestedness:* the observed weighted nestedness based on overlap and decreasing abundance (WNODA, green dot) is high compared with the distribution generated with the equiprobable null model, meaning that there is significant nestedness in the network. Nestedness is, though, very low compared to the distribution generated with the proportional null model, meaning that the network’s structure strongly deviates from a fully nested topology. We can, thus, conclude that there are important preferences in the pattern of observed interactions between plants and pollinators.

#### Binary nestedness

we analyzed the binary topology of the network using the NODF index. When weighted information was used to generate the proportional model, conclusions were similar to the analysis with the weighted index. However, when we based the proportional model on node degrees (marginal sums of the binary matrix) and connectance instead of total weights were conserved, using the null model 2 of Bascompte *et al*. (2003), it resulted in much lower values. In comparison, the observed nestedness (green dot) seemed higher than the expected given the degree distribution. This is, thus, a result of the distortion caused by using binary instead of weighted information. The distribution produced by the binary equiprobable null model was almost identical to the distribution produced by the weighted equiprobable null model and was not included in the figure.

## 4. DISCUSSION

### Nestedness and null models

In the present study we demonstrate that nestedness in a matrix can only result from marginal sum inequalities. This conclusion, arriving from a logical argument, corroborate previous studies that made similar claims based on comparisons between empirical networks and null models (Jonhson et al., 2013; Payrató-Borràs et al., 2019). We show that overlap, one of the two defining elements of nestedness, quantifies how well cell values are predictable by marginal sums. Preferences in the matrix, thus, reduce nestedness. In addition, for a matrix to present nestedness, it must have unequal marginal sums, otherwise marginal sums provide no information in addition to matrix dimensions and total sum.

This understanding delimits appropriate explanations for nestedness. The only way that a mechanism can generate nestedness is by increasing marginal sum inequalities, without adding preferences to the matrix. Thus, any explanation for nestedness in a species interaction network should address the question of why some species interact more frequently than others. Unequal abundances is an obvious first guess (Vázquez et al., 2007), but does not always apply (Vizentin-Bugoni et al., 2014). Other mechanisms may be involved, such as differences in fitness, activity, or detectability between species. Phylogenetic (Krasnov et al., 2012) and trait (Minoarivelo & Hui, 2016) specialization, on the other hand, lead to preferences and reduce the predictive power of second-order information on cell values, they thus reduce nestedness.

Analogously, in an archipelago, differences in island isolation and area may result in different chances of receiving or supporting species, generating a nested pattern of species occurrences. Local adaptation or competitive exclusion between pair of species, on the other hand, may result in preferences in the matrix, which decrease nestedness. Indeed, factors producing a hierarchy among sites or among species, and consequently skewed marginal sums, have been traditionally pointed out as the main causes of nestedness in species occurrences (Patterson & Atmar, 1986; Cutler, 1994).

Null model analysis has been a controversial step on nestedness tests, as there were no robust theory-oriented rules for choosing randomization algorithms, although this choice strongly affects the inference outcome (Ulrich et al., 2009). The new protocol proposed here overcomes this controversy by linking null models to explicit testable hypotheses. The equiprobable null model is used to test whether a given matrix present significantly higher nestedness than expected in the absence of nestedness-generating mechanisms. Matrices of an equiprobable null model thus present a random topology. The proportional null model tests whether the matrix is as nested as expected when nestedness-generating mechanisms predominate. Matrices of a proportional null model show a fully nested topology. This perspective challenges the perception that null model choice is a decision between type I and type II statistical errors: there is no common hypothesis being tested, but rather different hypotheses. Those null models are, thus, neither alternative nor conflicting, but can be used complementarily in nestedness analysis.

Ecological networks often show significant nestedness (Wright et al., 1997; Bascompte et al., 2003) and several previous studies looked for explanations for this high prevalence (e.g. Thebault & Fontaine, 2010). In our perspective, a high prevalence of nestedness is expected, as nestedness may easily arrive from many different ecological and evolutionary mechanisms (Valverde et al., 2018; Pinheiro et al., 2019). After all, unequal marginal sums promote nestedness (Krishna et al., 2008), and real-world networks are never formed by random connections between species with equal abundances and fitness. Furthermore, most species interaction networks in ecological literature represent taxonomically biased subsets, which often do not contain the strongest biological constraints and preferences observed in complete systems (Bezerra et al., 2009; Mello et al., 2011; Pinheiro et al., 2019).

Nevertheless, for a network to be as nested as expected by the proportional null model, it cannot include preferences between nodes. This may be the case in homogeneous and small-scale systems, but much less probable in more diverse networks (Lewinsohn et al., 2006). For instance, in a pollination network containing a few plants with similar flowers, we may expect no differential preferences of the pollinators for the plants. However, if the network comprises several plant families or plants with different pollination syndromes, it is more likely that different pollinators preferentially interact with each plant or group of plants (Cutler, 1994). Nested topologies are, thus, likely to occur in species interaction networks comprising low phylogenetic diversity (e.g. Flores *et al*., 2011), but unlikely to occur in diverse networks (e.g. Flores *et al*., 2013). In diverse networks, topologies that include preferences, such as modular and compound, tend to predominate (Mello et al., 2019).

### Methodological issues

Our study sheds light on the efficiency of our protocol in combination with nestedness indices. In weighted matrices, WNODA was very efficient in distinguishing between nested and random matrices. However, the use of binary information to produce proportional null models caused large distortions, both in our simulated matrices and in the case study, resulting in large underestimation of nestedness by the null model. This is not a trivial problem, as this kind of null model is the current standard methodology and has been widely applied for nestedness analysis in the past decades (Bascompte et al., 2003). Actually, it may be the cause of several studies finding higher nestedness in empirical binary ecological networks and metacommunities than in proportional null models (Patterson & Atmar, 1986; Bascompte et al., 2003). This diagnostic is corroborated by the absence of such pattern when quantitative information is used (Dormann et al., 2009; Joppa et al., 2010; Staniczenko et al., 2013) or when a more elaborate null model is applied (Payrató-Borràs et al., 2019). We echo previous authors in their call for a move beyond binary towards weighted matrices in ecological studies (Banašek-Richter et al., 2004; Scotti et al., 2007; Ings et al., 2009). We agree that there might be reasons to study the binary instead of the weighted structure of a matrix (Fründ et al., 2016). However, even in those cases, null models based on weighted information are more appropriate for nestedness analysis.

## Conclusion

Here we show that nestedness quantifies the effect of unequal marginal sums on the cell values of an interaction matrix. This change of perspective leads to a clearer understanding of nestedness and of the range of potential processes generating it. Moreover, by linking two different null models to explicit hypotheses, it promotes a more informative approach for testing nestedness, which go way beyond the standard dichotomic classification of networks as “nested or not”. We hope that this perspective contributes to solving old contradictions and paves the way towards a better understanding of the topology of species interactions, metacommunities, and other ecological and non-ecological systems.

## ACKNOWLEDGEMENTS

We thank our institutions and colleagues, who helped us in different ways during this project. The Ph.D. defense committee of RBPP, composed by Tatiana Cornelissen, Aristóteles Neto, Mario Almeida-Neto, and Paulo Peixoto, provided suggestions that considerably improved the quality of our study. The Graduate School in Ecology of the Federal University of Minas Gerais (ECMVS), Brazil, provided RBPP with a Ph.D. scholarship from the Brazilian Council for Scientific and Technological Development (CNPq). Infrastructure for this study was provided by ECMVS, the Department of Ecology of the University of São Paulo (USP), Brazil, and the Department of Biometry and Environmental System Analysis of the University of Freiburg, Germany. The Graduate School in Ecology of the State University of Campinas (PPGE-UNICAMP), Brazil, provided GMFF with a scholarship from the Brazilian Coordination for the Improvement of Higher Education Personnel (CAPES). RBPP received a scholarship from the joint program between CAPES, CNPq, and Deutscher Akademischer Austauschdienst (DAAD) (88887.161398/2017-00) and is currently funded by São Paulo Research Foundation (FAPESP, post-doctoral grant 2020/06771-2). MARM was funded by the Minas Gerais Research Foundation (FAPEMIG: PPM-00324-15), Alexander von Humboldt Foundation (AvH, 3.4-8151/15037 and 3.2-BRA/1134644), CNPq (302700/2016-1 and 304498/2019-0), Dean of Research of the University of São Paulo (PRP-USP, 18.1.660.41.7), and FAPESP (2018/20695-7).

## AUTHORS’ CONTIBUTIONS

All authors conceived the ideas and designed the study. RBPP led the analysis and the writing of the manuscript, with help from all authors.

## DATA AVAILABILITY

The source of the dataset used as a case study is referenced in the text. Commented scripts to reproduce the study and functions for applying the proposed procedures on alternative networks are available in GitHub (https://github.com/pinheirorbp/nestedness).

## Notes

https://github.com/pinheirorbp/nestedness

### Competing Interest Statement

The authors have declared no competing interest.

