## Appendix S1 for "Linking null models to hypotheses to improve nestedness analysis"

### Appendix S1. Efficiency analysis

Pinheiro, R.B.P., Dormann,C.F., Felix, G.M.F, and Mello, M.A.R.

November 27, 2021

#### WEIGHTED MATRICES

In our novel perspective, the main objective of a nestedness test is to distinguish between matrices with randomly distributed cell values (non-significant nestedness, equiprobable model), matrices in which cell values are partially defined by the marginal sums (significant nestedness), and matrices in which cell values are fully determined by marginal sums (nested matrices, proportional model). Here, we analyzed the capacity of several indices in distinguishing these topologies following our protocol.

We produced probability matrices with dimensions: 5x5, 10x10, and 20x20, based on three different marginal probabilities: lognormal, linear decrease, and equiprobable. Then, we generated matrices from these probability matrices with different total samplings: 50, 100, 200, 400, 800, 1600, 3200, and 6400. For each unique setup we produced 10,000 matrices. For each matrix we calculated NODF, WNODF and WNODA.

In this analysis we used both weighted (WNODF and WNODA) and binary (NODF) indices. However, the models were always produced using the weighted information, and indices are compared between matrices with fixed sampling (instead of connectance).

Here we present plots for a graphycal evaluation of the efficiency to separate between the proportional (lognormal and linear decrease) and the equiprobable models. Indices in the y-axis and sampling (total weights on the matrix) on the x-axis. Each plot present median and intervals containing 95% of points. NODF, WNODF and WNODA values were divided by 100.

**MATRIX SIZE: 20 x 20**

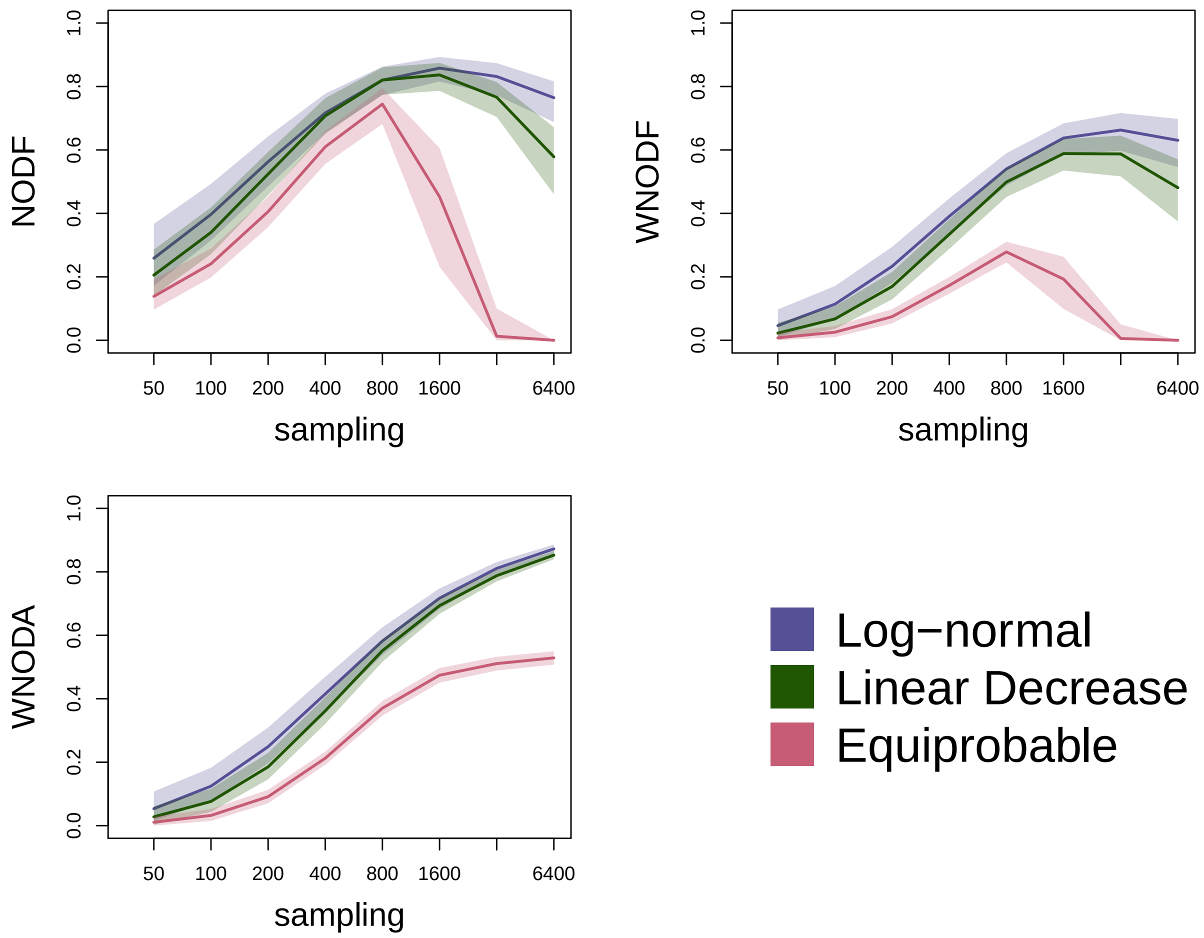

**MATRIX SIZE: 10 x 10**

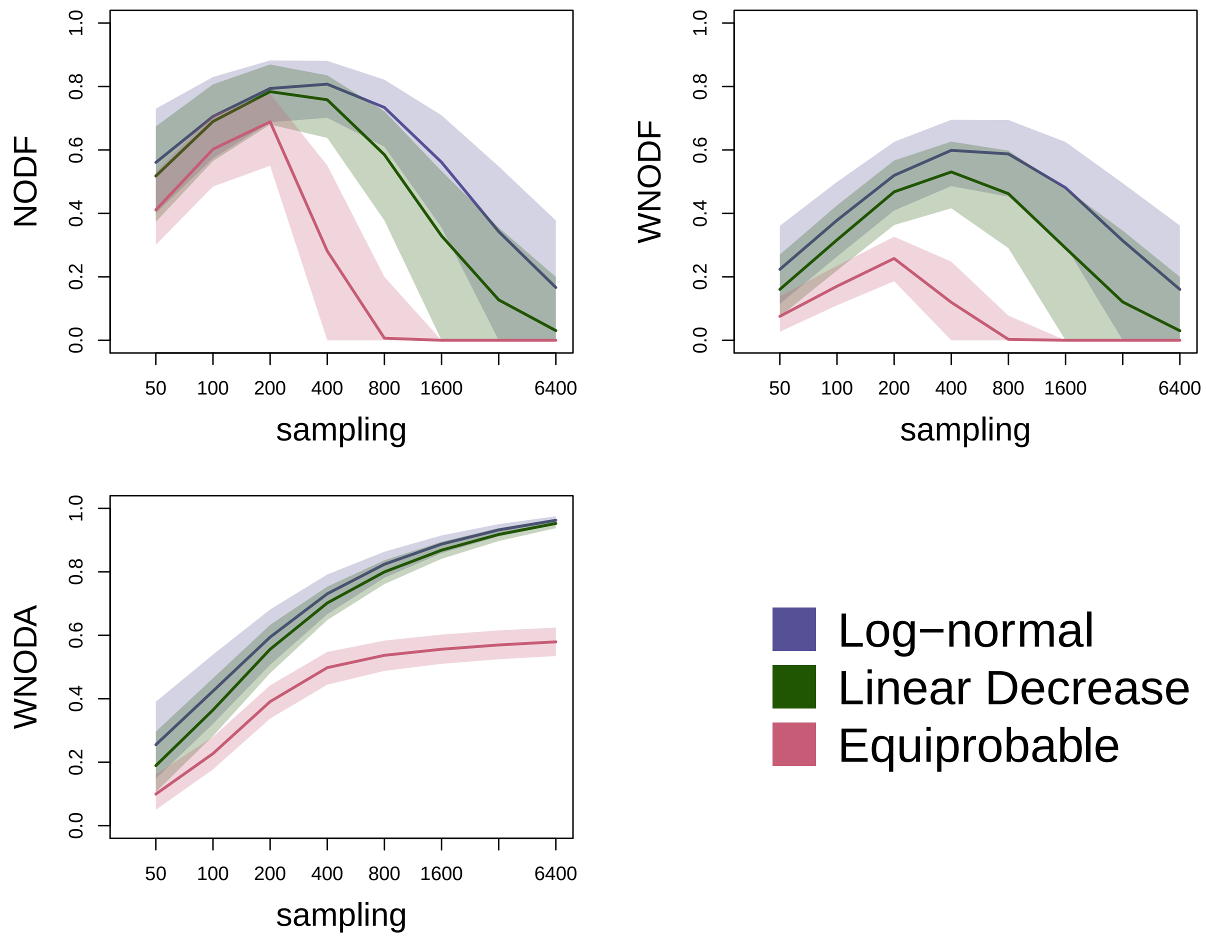

**MATRIX SIZE: 5 x 5**

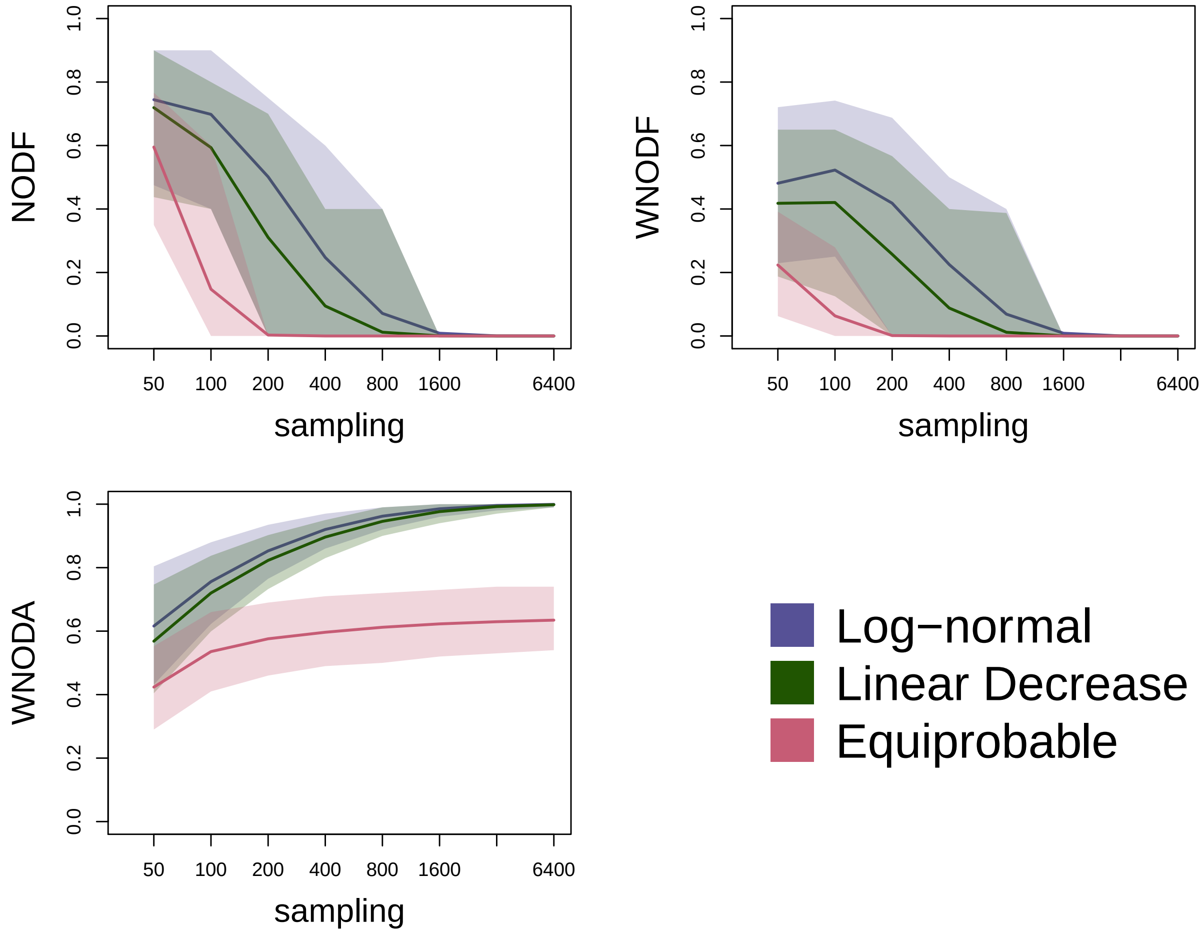

### BINARY MATRICES

In our novel perspective, the main function of a nestedness test is to distinguish between matrices with randomly distributed cell values (non-significant nestedness, equiprobable model), matrices in which cell values are partially defined by the marginal sums (significant nestedness), and matrices in which cell values are fully determined by marginal sums (nested matrices, proportional model). Here, we analyzed the capacity of NODF in distinguishing these topologies following our protocol.

We produced probability matrices with dimensions: 5x5, 10x10, and 20x20, based on three different marginal probabilities: lognormal, linear decrease, and equiprobable. In Appendix S1 we generated weighted matrices from these probability matrices with fixed total sampling. Here, we produced binary matrices for a more appropriate analysis of NODF, as a binary index.

As binary matrices do not present weighted information we can only fix the connectance. We produced binary matrices with connectances: 0.3, 0.5, 0.7, 0.9, 0.92, 0.94, 0.96, and 0.98. For each matrix we calculated NODF.

Moreover, in analysis of binary matrices, we cannot know the original marginal probabilities, only the binary marginal sums (node degrees). To inspect the distortion caused by this approximation, for each matrix produced with fixed connectance, we produced a randomized matrix using a proportional algorithm based on binary marginal sums (binary proportional).

Here we present plots for a graphycal evaluation of the capacity to separate between the proportional (lognormal and linear decrease) and the equiprobable null models. We also present values of the binary proportional null model. Indices in the y-axis and connectance on the x-axis. Each plot present median and intervals containing 95% of points. NODF values were divided by 100.

**MATRIX SIZE: 20 x 20**

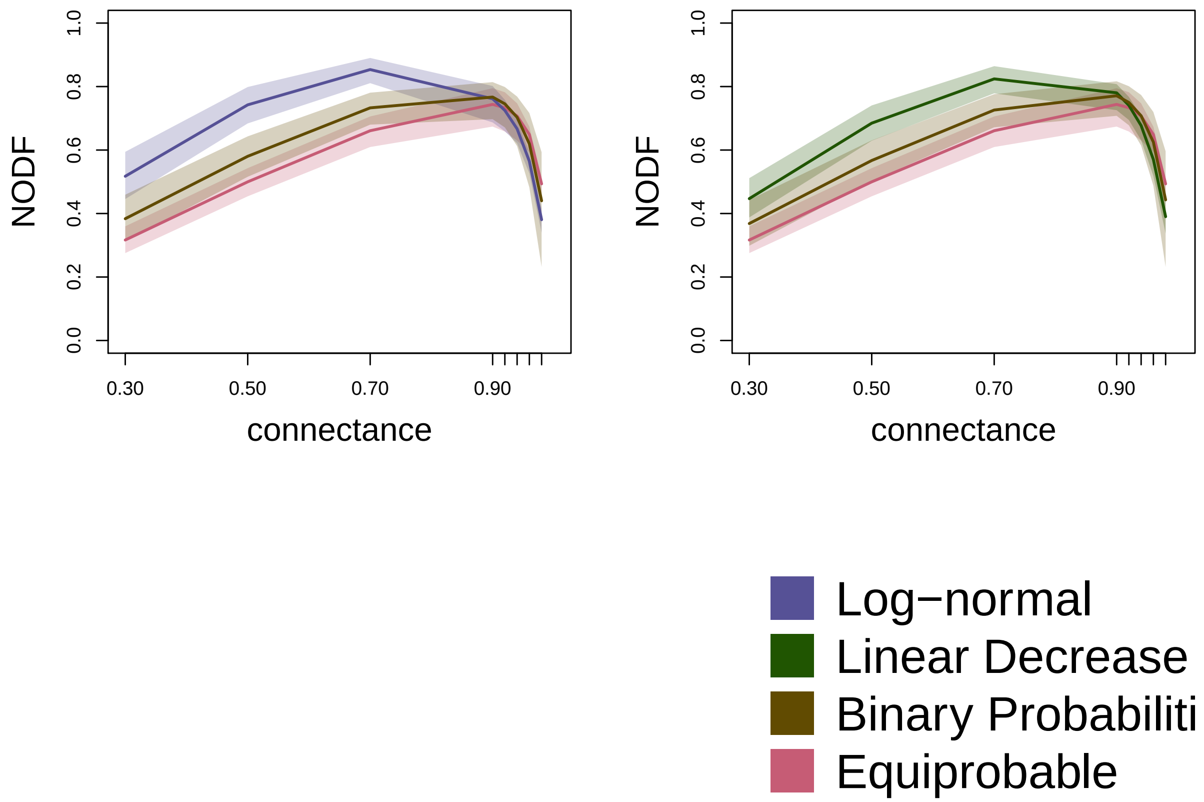

**MATRIX SIZE: 10 x 10**

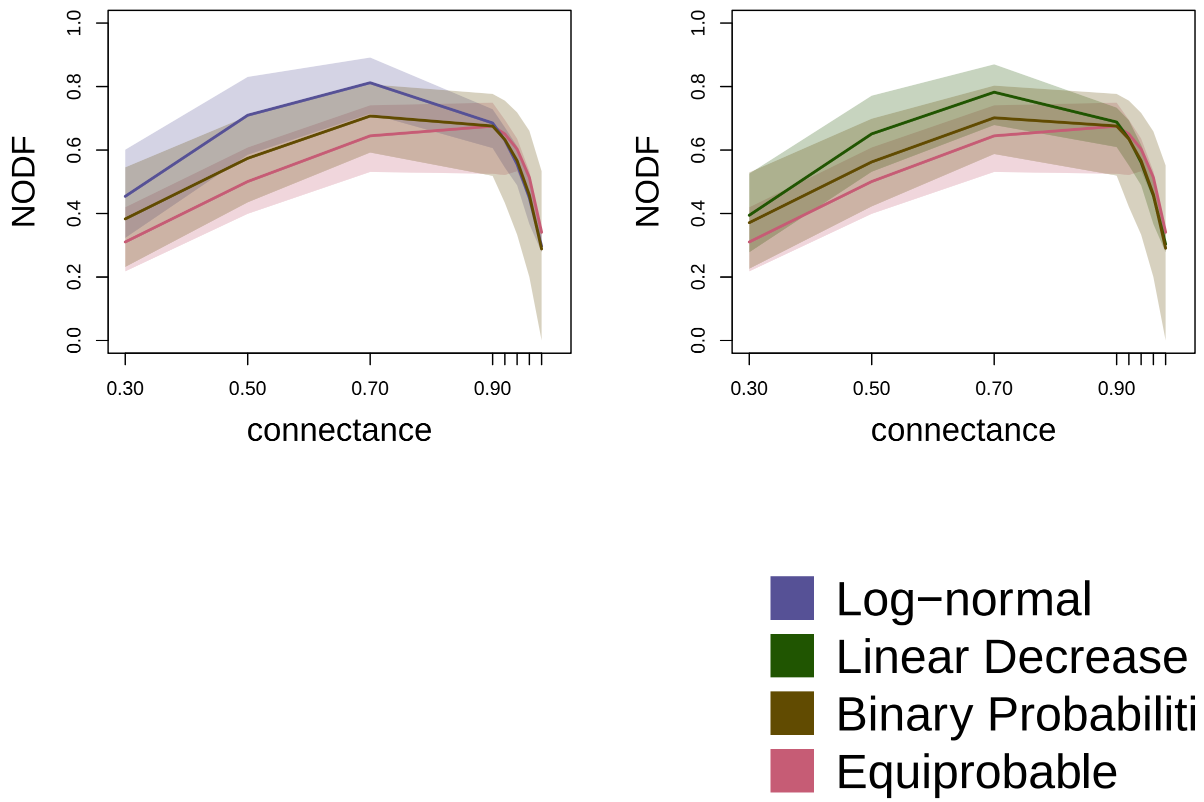

**MATRIX SIZE: 5 x 5**

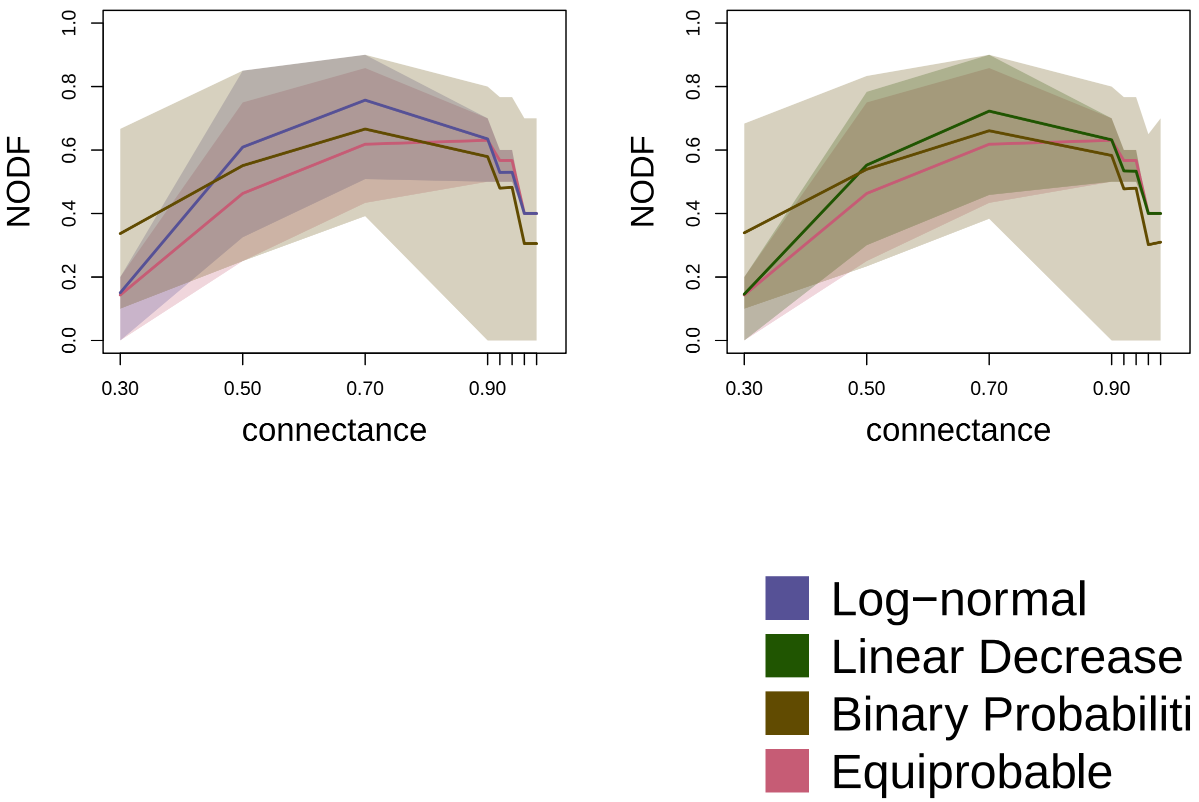
