## Supplementary material for "Linking null models to hypotheses to improve nestedness analysis": Figures S1 - S2

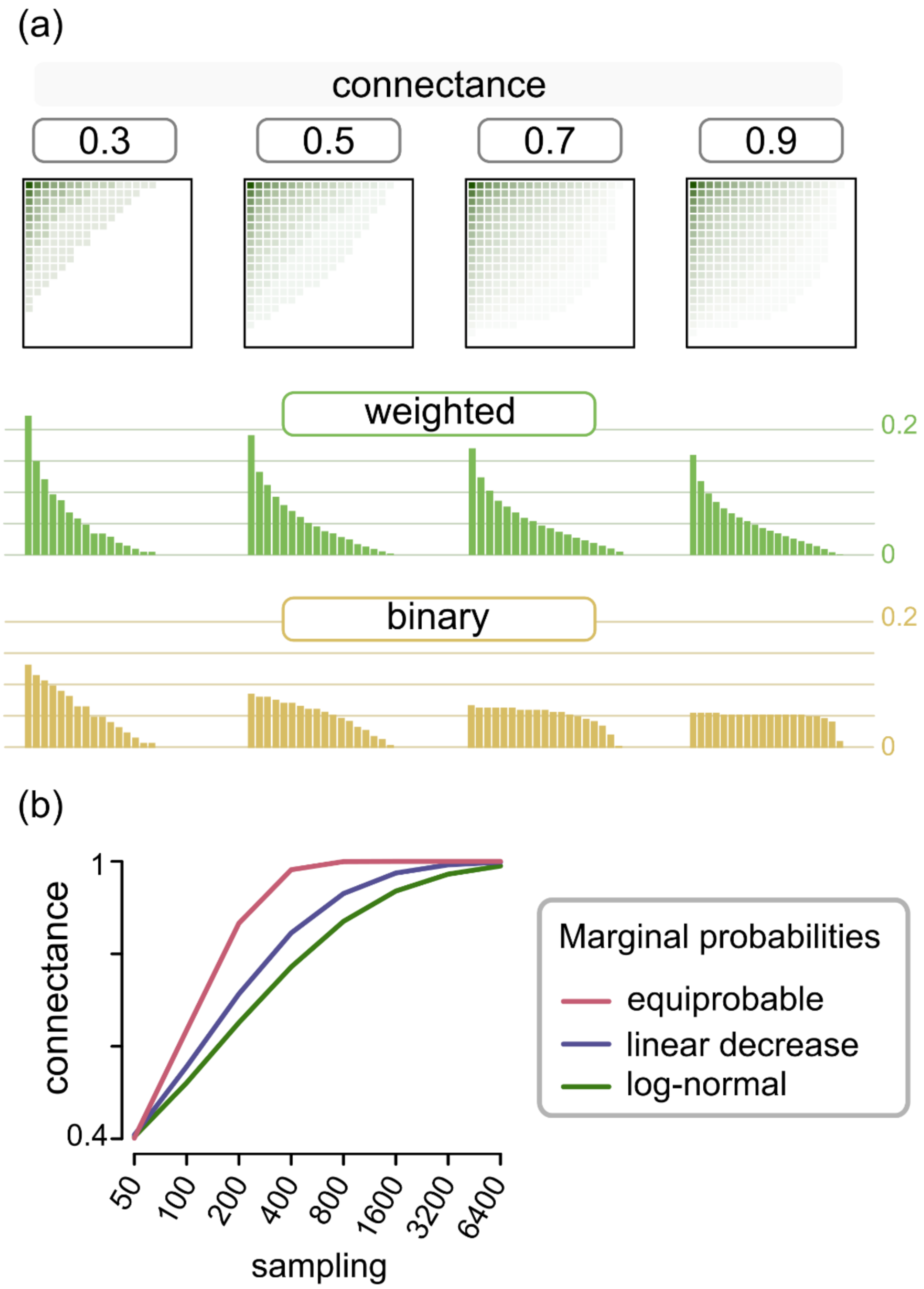


**FIGURE S1** (a) Binary marginal sum distributions are contingent on matrix dimensions and connectance. They are, therefore, poor proxies for weighted marginal sum distributions, especially in highly connected matrices. Matrices were produced following a log-normal distribution of marginal sums. Relative marginal sums are represented as bars. For simplicity we did not remove the empty rows and columns of the matrices, which would modify the connectance (actual scores are 0.47, 0.62, 0.78, and 0.9). (b) Connectance is not a direct proxy for the first-order information on the weighted matrix. We created probability matrices (dimensions: 10 x 10) based on marginal probability distributions and sampled interactions, to produce interaction matrices. For each marginal probability we created 10,000 matrices and measured their connectance at different sampling intensities. In the plot we see that the mean connectance (lines) is not only affected by sampling, a first-order information, but also by marginal probability distribution, a second-order information of the weighted network.

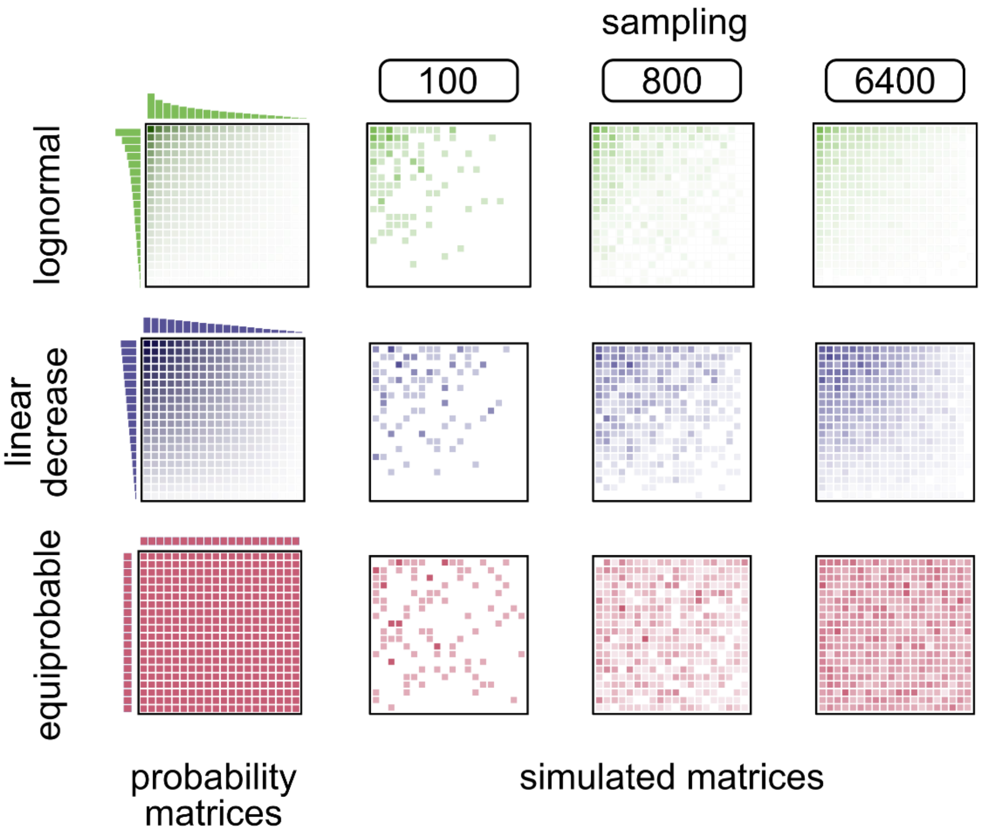


**FIGURE S2** To test whether nestedness indices are efficient in distinguishing matrices with different topologies under our protocol, we simulated matrices following different probability distributions. First, we produced probability matrices with three different distributions of node probabilities: log-normal, linear decrease, and equiprobable. Then, we generated randomized matrices with different total sums (sampling) following cell probabilities.
